# Transient amplitude modulation of alpha-band oscillations by short-time intermittent closed-loop tACS

**DOI:** 10.1101/2020.01.23.916676

**Authors:** G. Zarubin, C. Gundlach, V. Nikulin, A. Villringer, M. Bogdan

**Affiliations:** Technical Informatics Department, Leipzig University, Leipzig, Germany; Department of Neurology, Max Planck Institute for Human Cognitive and Brain Sciences, Leipzig, Germany; Institute of Psychology, University of Leipzig, Leipzig, Germany; MindBrainBody Institute at the Berlin School of Mind and Brain, Humboldt-Universität zu Berlin, Berlin, Germany; Neurophysics Group, Department of Neurology, Campus Benjamin Franklin, Charité Universitätsmedizin Berlin, Berlin, Germany; Department of Cognitive Neurology, University Hospital Leipzig, Leipzig, Germany; Center for Stroke Research Berlin, Charité – Universitätsmedizin Berlin, Berlin, Germany; Centre for Cognition and Decision Making, National Research University Higher School of Economics, Moscow, Russian Federation

**Author notes:** Co-first authors (G. Zarubin, C. Gundlach).

## Abstract

Non-invasive brain stimulation techniques such as transcranial alternating current stimulation (tACS) have recently become extensively utilized due to their potential to modulate ongoing neuronal oscillatory activity and consequently to induce cortical plasticity relevant for various cognitive functions. However, the neurophysiological basis for stimulation effects as well as their inter-individual differences are not yet understood. In the present study we used a closed-loop EEG transcranial alternating current stimulation protocol (EEG-tACS) to examine the modulation of alpha oscillations generated in occipito-parietal areas. In particular, we investigated the effects of a repeated short-time intermittent stimulation protocol (1 s in every trial) applied over the visual cortex (Cz and Oz) and adjusted according to the phase and frequency of visual alpha oscillations on the amplitude of these oscillations. Based on previous findings, we expected higher increases in alpha amplitudes for tACS applied in-phase with ongoing oscillations as compared to an application in anti-phase and this modulation to be present in low-alpha amplitude states of the visual system (eyes opened) but not high (eyes closed).

Contrary to our expectations, we found a transient suppression of alpha power in inter-individually derived spatially specific parieto-occipital components obtained via the estimation of spatial filters by using the common spatial patterns approach. The amplitude modulation was independent of the phase relationship between tACS signal and alpha oscillations, and the state of the visual system manipulated via closed- and open-eye conditions. It was also absent in conventionally analyzed single-channel and multi-channel data from an average parieto-occipital region.

The fact that the tACS modulation of oscillations was phase-independent suggests that mechanisms driving the effects of tACS may not be explained by entrainment alone, but rather require neuroplastic changes or transient disruption of neural oscillations. Our study also supports the notion that the response to tACS is subject specific, where the modulatory effects are shaped by the interplay between the stimulation and different alpha generators. This favors stimulation protocols as well as analysis regimes exploiting inter-individual differences, such as spatial filters to reveal otherwise hidden stimulation effects and, thereby, comprehensively induce and study the effects and underlying mechanisms of tACS.

## Introduction

Non-invasive brain stimulation (NIBS) technology has gained increasing attention in the last few years from the scientific community (Bergmann et al., 2016; Thut et al., 2017; Antal et al., 2017; Vosskuhl et al., 2018), clinical (Yavari et al., 2017; Palm et al., 2014), sports (Edwards et al., 2017; Angius et al., 2018), military (Nelson et al., 2016), and other fields. One of the reasons for this growing interest is the successful modulation of cognitive, motor, and perceptual functions in numerous studies in different domains such as motor function (Brittain et al., 2013; Feurra et al., 2011; Angius et al., 2018), visual (Helfrich et al., 2014; Zaehle et al., 2010), auditory (Riecke et al., 2015), somatosensory (Feurra et al., 2011; Gundlach et al., 2016, 2017), or linguistic processing (Riecke et al., 2018; Wilsch et al., 2018) and for higher cognitive functions such as decision making, creativity, or self-aware dreaming (Lustenberger et al., 2015; Sela et al., 2012; Voss et al., 2014). Another reason is the widespread availability of various experimental, clinical protocols and instructions (Bergmann et al., 2016; Antal et al., 2017; Tavakoli & Yun, 2017). In addition, NIBS is generally a safe and well tolerated form of brain stimulation with a comparatively simple set up. Among the broad variety of existing NIBS techniques, transcranial magnetic and direct current stimulation (TMS, TDCS) are the most widely used, and mechanisms of their actions and induced effects seem to be the best understood (Pelletier & Cicchetti, 2015; Chervyakov et al., 2015). However, transcranial alternating current stimulation (tACS) also has broad applications due to its ability to modulate ongoing neural oscillatory activity in a flexible way by precisely tuning stimulation parameters (such as frequency, phase, amplitude, or a combination of these) to each individual or each experimental session (Herrmann et al., 2013; Reato et al., 2013).

The advantages of tACS may be exploited even further in case of an adaptive or closed-loop approach, when stimulation parameters are tuned online during the experiment in a particular determined manner (Karabanov et al., 2016; Zrenner et al., 2016). In such a framework, brain responses to the stimulation, usually obtained from electroencephalography (EEG) or magnetoencephalography (MEG) data, serves as feedback and is used for the modification of control parameters. Significant efforts in recent research have been devoted to the establishment and application of closed-loop tACS-EEG/MEG models (Bergmann et al., 2016; Thut et al., 2017). However, despite an increasing number of proposed models, the field of adaptive tACS still lacks experimentally validated solutions. This can be explained by the complexity of the task: implementation and utilization of closed-loop tACS has various technical challenges and fundamental questions, which narrow and delay the development of this field.

First of all, despite numerous studies, the exact neural mechanisms of the effects of tACS are still not well understood. Animal as well as human and computational modeling studies suggest that weak alternating electric fields modulate spiking patterns of neurons by means of neural entrainment (Deans et al., 2007; Fröhlich & McCormick, 2010; Ozen et al., 2010; Helfrich et al., 2014) or induction of spike-timing plasticity (Zaehle et al., 2010; Polanía et al., 2012; Vossen et al., 2015; Silva et al., 2018). As suggested by resonance theory, these effects should be highly frequency dependent and observable online during stimulation (Hutcheon & Yarom, 2000). In line with this, various studies have reported online effects of tACS on behavior (Kanai et al., 2010; Vosskuhl et al., 2015; Strüber et al., 2013; Gundlach et al., 2016) as well as markers of neural activity in the EEG, MEG or fMRI (Witkowski et al., 2015; Neuling et al., 2015; Cabral-Calderin et al., 2015). However, it has recently been shown (Asamoah et al., 2019) that effects of tACS on the motor system are, at least, partly related to transcutaneous stimulation of peripheral nerves in the skin beyond transcranial stimulation of cortical neurons (however, see Krause et al., 2019, Vieira et al., 2019). Examining stimulation mechanisms therefore remains an open question. Similar to the logic of applying tDCS, research questions concerning the application of tACS often focus on the offline effects of stimulation after a prolonged application. As suggested in some studies, processes leading to online effects of tACS and those leading to offline effects that sustain (or even manifest) after stimulation may rely on different neural mechanisms (Reato et al., 2010; Strüber et al., 2015). In a recent review paper on the immediate effects of tACS (Liu et al., 2018), five possible neural mechanisms were suggested: “stochastic resonance, rhythm resonance, temporal biasing of neuronal spikes, entrainment of network patterns, and imposed patterns”. Importantly, how these mechanisms contribute to observed effects specifically and how they interact or compensate each other is largely unknown.

Secondly, one of the biggest challenges with tACS and NIBS in general is high inter- and intra-individual variability of effects (Ziemann et al., 2015). As mentioned by Guerra et al. 2017, this variability can be due to different factors, including physiological (variability of brain morphology, endogenous states, and different responses to stimulation), technical (particular setup and parameters of stimulation), and also statistical differences (numbers of participants and trials per groups and conditions).

Thirdly, the main technical challenge for analyzing and investigating online effects during stimulation is the induction of stimulation artifacts, which exceed the measured neural signals by several magnitudes. Although there is an increasing number of propositions for overcoming this challenge and some prominent solutions (Witkowski et al., 2015; Kohli & Casson, 2019; Kasten et al., 2018; Noury & Siegel, 2017) directed to elimination of artifacts, such approaches still require sophisticated experimental designs and/or heavy computational procedures. Most human tACS studies have used stimulation protocols with a duration applied in the range of minutes. However, the minimum duration of stimulation required for inducing after-effects, such as a modulation of coherence or power of oscillations, has not been established. Previous studies using conventional tACS (Vossen et al., 2015), found that intermittent tACS in the alpha range, applied over the visual cortex with trains of 8 s, but not 3 s, led to a modulation of visual alpha amplitude. Strüber et al. (2015) also applied tACS intermittently with 1 s intervals, but found no significant after-effects. Because the effects of tACS are proposed to arise from an interaction of the applied electromagnetic field and ongoing neural oscillations, the state of neural activity before and during stimulation is assumed to be an important factor for any stimulation outcome (Feurra et al., 2013; Neuling et al., 2013). Therefore, the relation between parameters such as phase, amplitude, and frequency of the targeted neural oscillations and applied tACS signal should modulate the effects of tACS. In line with resonance theory leading to an online entrainment of ongoing oscillations by tACS, the impact of these parameters may be more pronounced for shorter stimulation intervals (i.e., for longer stimulation intervals initial differences between the tACS phase and the phase of ongoing alpha oscillations, for instance, may be neglectable, as the phase synchronization should obtain a maximum after some time). The application of tACS phase-synchronized with ongoing alpha oscillations follows this logic, where a closer coupling between tACS and alpha-band oscillations should enhance potential stimulation effects and render tACS effective, even for low stimulation durations previously found ineffective.

In this work, we present results of a study using a closed-loop EEG-tACS protocol by which the stimulation signal was phase coupled with ongoing alpha oscillations. Our research is based on a closed-loop model that we developed and presented previously (Zarubin et al., 2018), using an approach with intermittent stimulation. In particular, we investigated the effects of repeated short durations of tACS (1 s) applied over the visual cortex on alpha amplitude when tACS signals and ongoing alpha visual alpha oscillations were phase-synchronized (in-phase) or in opposite phase (anti-phase) and in different states, when alpha levels were high in amplitude (eyes closed) or low (eyes opened). We expect higher increases in alpha amplitudes for tACS applied in-phase as compared to an application in anti-phase and this modulation to be present in low-alpha amplitude states of the visual system (eyes opened), but not high (eyes closed). A dependency of stimulation effects on the phase of the tACS signal would also examine online entrainment of visual alpha oscillations by tACS as a candidate mechanism for tACS effects. In addition to tailoring tACS to each subject’s individual alpha frequency and phase, we also adapted the analysis regime to acknowledge inter-individual differences in the spatial pattern of stimulation effects. Specifically, we studied potential modulations of neural oscillations conventionally, using single-channel and multi-channel data, from the parietal-occipital region and contrasted it with data individually derived via the application of spatial filters with the common spatial patterns approach (Blankertz et al., 2008).

## Materials and Methods

### Participants

Twenty healthy adults (9 females, mean age 28.4 ± 3.2 years) took part in this experiment and received monetary compensation for their participation. None of the participants had a history of psychiatric or neurological diseases and none were on any current medication affecting the central nervous system. Participants were informed about all aspects of the study and gave their written informed consent prior to the experiment. The study protocol was approved by the local ethics committee (“Modulation neuronaler Oszillationen mittels transkranieller Wechselstromstimulation und ihr Effekt auf die somatosensorische Wahrnehmung”, 12.08.2014, Reference number: 218-14-14072014). None of the participants have claimed that the feeling of stimulation was unpleasant, and none of them have experienced phosphenes.

### Experimental procedure

Each experimental session consisted of a preparation and information part, as well as the actual stimulation and EEG recording. During the preparation and information part, participants were introduced to the aim, set-up, and procedure of the study as well as technical background of the stimulation. Individual contraindications for transcranial alternating current stimulation were checked and the consent form was given, explained, and signed. tACS electrodes as well as EEG electrodes were then applied and set up. Participants were subsequently instructed to sit relaxed, avoid movements, and later to keep their eyes opened or closed depending on the stimulation block. The experiment included one session, which was completed by all participants. At the beginning, four 1-min resting state EEG data (2 eyes closed [EC], 2 eyes opened [EO], one after another) were collected to determine individual alpha frequencies (IAFs) by contrasting peaks in FFT spectra between EC and EO states at channel POz. Approximately five short stimulation sequences were then applied to test correct functioning of closed-loop model units. Thereafter, 10 blocks of 50 trials were executed consecutively, starting with either the EC or EO state, following one after another with short breaks between the blocks. Each trial consisted of a 1 s pre-stimulation interval, 1 s of stimulation (with phase adjusted by prediction from pre-stimulation), a 1 s post-stimulation interval, and an inter-trial interval (ITI). The random ITI was in the range of 333 to 666 ms with a mean value of 500 ms. Every block consisted of 25 in-phase and 25 anti-phase stimulation trials in randomly shuffled order (Figure 1b). The total time for each block was 3 min, resulting in a total time of approximately 45 min for all 10 blocks with breaks between and a total stimulation time of 8 min, 20 s. Our study design thus contained the factors STATE (eyes open vs. eyes closed) and STIMULATION (in-phase vs. anti-phase).

**Figure 1.**
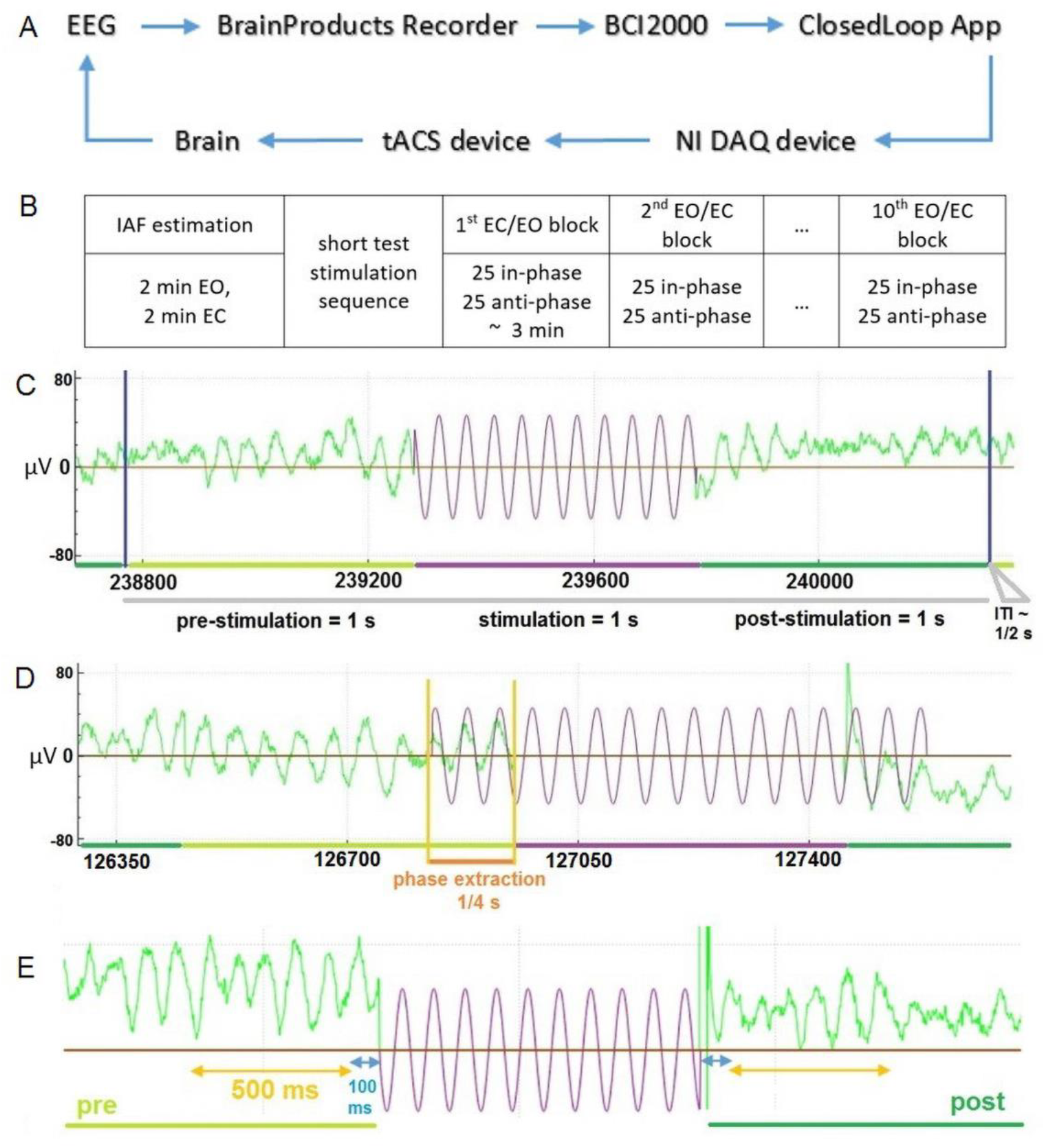
General description of the Closed-Loop-Model and the temporal structure of the data A) The general model scheme of the Closed-Loop setup B) Outline of experimental procedure C) An exemplary temporal trial structure with data in green representing recorded EEG data and data in violet depicting stimulation signal D) Illustration of the stimulation signal estimation based on an extraction interval of 0.25 s for an exemplary “in-phase” stimulation trial E) Temporal scheme of EEG data (green) used for analysis of stimulation effects due to tACS (in violet – schematic representation of stimulation wave). Interval in blue represents 100 ms data interval excluded in order to reduced stimulation artifacts and potential edge filtering effect

### Electrical Stimulation

tACS electrodes (two conductive rubber 4 × 4 cm) were attached over standardized Cz and Oz channel locations (Herwig et al., 2003; Jasper, 1958) underneath the EEG recording cap, and sinusoidal alternating current was applied at IAF (calculated according to the procedure described above) using a battery-driven stimulator (DC-Stimulator Plus, NeuroConn, Ilmenau, Germany). Stimulation electrode positions were selected based on previous studies, in which modulations of visual alpha oscillations by tACS were reported (Neuling et al., 2013; Helfrich et al., 2014; Ruhnau et al., 2016). Impedances were kept below 10 kΩ with Ten20 conductive paste (Weaver and company, Aurora, Colorado). tACS at IAF was applied with an intensity of 1 mA (peak-to-peak) for all subjects. The stimulation signal for every trial was initially determined in a closed loop application (custom made, C++), then generated through NI DAQ card (USB 6343, National Instruments, Texas, USA) and transmitted to the DC-Stimulator Plus “remote input” port.

### EEG recording

EEG data was recorded using Brain Products amplifier BrainAmp MRplus (Brain Products GmbH, Gilching, Germany) with 31 Ag-AgCl electrodes mounted in an passive EEG EasyCap using a standard 10–20 system layout without Oz and Cz electrodes, with reference and ground electrode positioned at FCz and AFz and applying a sampling rate of 500 Hz. The low sampling rate was used to reduce data transfer delays through components of the model, and during recording the data was streamed from BrainVision Recorder through its RDA client to BCI2000 (open-source software, SchalkLab), which then transmitted it to the ClosedLoop application for analysis and optimal phase prediction.

### Closed-loop model

The general model scheme is represented in Figure 1A. The model is implemented with 3 modes of functioning: “Online”, “Offline”, “Record-only” / “Stimulation-only”. In “Online” mode (Figure 1C) the system functions in a state required for adaptive stimulation with short interval cycles (e.g., 1 s) and incorporates: import of EEG signals (pre-stimulation interval), estimation of optimal parameters (e.g., phase shift) and calculation of stimulation signal, computation of required optimization time, compensation of optimization and transduction delays, and transmission of stimulation signal (stimulation interval) through a NI DAQ card to the stimulator device input (BNC port). A typical cycle also contains a post-stimulation interval for analysis of effects and inter-trial interval. The “Offline” mode was used for the testing of phase prediction methods, while the “Record-only” and “Stimulation-only” modes can be used for continual recording of data and for continual stimulation with fixed parameters. All computations were performed on Lenovo P70 (Intel i7 OctaCore 2.6GHz, 16Gb RAM, Lenovo Group Limited, Beijing, China).

### Phase prediction, delay estimation and compensation

For phase prediction we utilized the same Hilbert-based approach as tested and validated in our previous study (Zarubin et al., 2018). This method is based on the assumption that with short time intervals we can consider phase dynamics to be quasi stationary, which allows us to perform phase prediction by the extraction phase information from a current (pre-stimulation) interval in order to forecast phase dependent stimulation for the following interval. Importantly, we only used a predefined part of the pre-stimulation interval for phase extraction (extraction interval), which is controlled by a parameter and was settled to the last quarter (250 ms) of the whole interval (Figure 1D). For the extraction interval, phase value should be optimal to achieve the best prediction. We used Hilbert transformation (after FIR filtering, 100 ms length) to obtain instantaneous phase values and iterative search across sine waves with different phases for this optimization. Euclidean difference (L^2^ norm) for vectors of instantaneous phase between extraction interval and various generated sine waves was criteria for minimization -in case of “in-phase” relation, and maximization in case of “anti-phase” relation. Transduction and optimization delays were estimated and compensated with the same procedure as validated in (Zarubin et al., 2018). For the chosen phase prediction method (Hilbert-based), the average value of optimization delay across all subjects was 15±2.4 ms and average transduction delay was 72 ms.

### Time intervals

We considered time windows of 500 ms in length (see Figure 1E), extracted from pre- and post-simulation intervals. We omitted the first 100 ms from the beginning of the post-stimulation interval, and the last 100 ms from the end of pre-stimulation. This reduction was applied to avoid influence of the filtering edge effect and possible stimulation artifacts on further analysis. Moreover, we examined the temporal evolution of the stimulation effect by testing potential differences of four different 500-ms long overlapping post-stimulation time windows ([100; 600] ms, [200; 700] ms, [300; 800] ms, [400; 900] ms in relation to onset of the stimulation) as compared to the same pre-stimulation time window ([−600; −100] ms).

### Common Spatial Patterns

The motivation for the application of spatial filters to investigate effects of stimulation is driven by the fact that while effects of tACS have a relatively broad distribution, they still influence a rather well-defined neuronal population. The extraction of neural activity from such a population, however, is hampered by volume conduction, which leads to the EEG recording reflecting many neuronal populations with overlapping fields. Thus, in order to extract the EEG activity most susceptible to stimulation, we use spatial filtering that maximizes/minimizes the difference in power between the reference and post-stimulus intervals. We can accomplish this using spatial filters that maximally discriminate between on and off stimulation intervals. One of the established and efficient methods for extraction of modulated brain activity is the Common Spatial Patterns (CSP) approach, which was introduced in (Blankertz et al., 2008) for single-trial analysis and is commonly used in brain-computer interface (BCI) applications to differentiate between classes of brain activation. Moreover, CSP has been successfully applied previously for the extraction of alpha band activity sensitive to the processing and discrimination of standard and deviant visual stimuli (Tugin et al., 2016).

We utilized CSP to obtain data from parietal-occipital regions to investigate possible differences between pre- and post-stimulation activity for alpha band oscillations in visuo-occipital components. Firstly, channels affected by sustained electrical current from neighboring tACS electrodes were rejected via EEGLAB functions and visual inspection. Then we extracted and merged all remaining data from pre- and post-stimulation intervals in two matrices (S_pre_,S_post_). The post-stimulation data collection began 80 ms after stimulation offset, the arbitrary 80 ms shift was applied to avoid possible stimulation-related artifacts. Further, the data intervals were band-pass filtered with a frequency range of 7 to 14 Hz with 4th order Butterworth zero-phase filter to prevent edge effects on borders caused by baseline shifts during experiment. Next, we constructed two new matrices (S’_pre_ and S’_post_) based on S_pre_ and S_post_ by cutting respectively the last and first 100 ms from each interval, of the original matrices, to avoid the influence of filtering edge effects would have on further analysis. Then, following the scheme described earlier (*Time intervals*, Figure 1e), 500 ms of data preceding the stimulation was used as the pre-stimulation interval and 500 ms following the stimulation as the post-stimulation data. Further, covariance matrixes (*C*_*pre*_,*C*_*post*_) for pre- and post-stimulation states were determined (1):

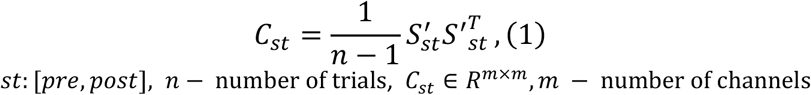

Afterwards generalized eigenvalue decomposition of (*C*_*pre*_,*C*_*pre*_ + *C*_*post*_) was computed (2):

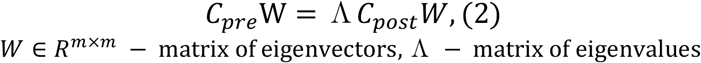

Eigenvectors *w*_*j*_ (*j* = 1: *m*) of the decomposition are CSP filters, where eigenvectors corresponding to higher eigenvalues maximize pre-stimulation activity (i.e. a decrease in alpha power from pre- to post-stimulation period). Eigenvectors with the lower eigenvalues maximize post-stimulation activity (i.e. an increase in alpha power from pre- to post-stimulation period). Vectors of the inverse matrix *W*^−1^ represent spatial patterns. When the CSP filters were computed, for every subject from the same order of filters we selected one filter maximizing pre-stimulation alpha power over post-stimulation power (CSP(pre): *w*_*j*_) and one maximizing post- over pre-stimulation alpha power (CSP(post): *w*_*m*−*j*+1_). Selection of order *j* was performed by analyzing the topographies of the CSP filters (with the inverse matrix *W*^−1^ and the *topoplot()* function in EEGLAB). Here the first filter was chosen for which the spatial pattern showed a parietal-occipital topography. This allowed us to consider specifically alpha oscillations from regions targeted with tACS. Importantly, to avoid overfitting, we implemented a cross validation procedure for every trial. For each single trial, spatial filters were first constructed based on data, not from that trial, but from all other trials (i.e., a leave-one-out procedure). Then, the selected CSP filter was normalized by its standard deviation (3) and applied to the trial to obtain the projected component *P*_*pre*/*post,i*_ of a current trial *X*_*i*_ (4):

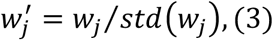

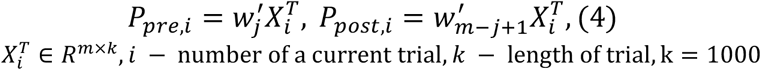

Crucially, the CSP estimation and selection was thus independent of the data for which these filters were then adopted. Finally, the obtained component *P*_*pre,i*_ of all trials were merged and used in the further analysis as CSP(pre) data, whereas the components *P*_*post,i*_ were merged and used as CSP(post) data.

### Source channel, parietal-occipital channels, and CSP components

Data from two sets of CSP components were compared to a more conventional data selection procedure. More specifically, CSP(pre) maximized alpha power “pre over post” while CSP(post) maximized alpha power “post over pre”. Conventional data selection was either based on data from a single channel (POz; closest to stimulation electrode) or from a parietal-occipital cluster (POC-region) with nine channels (P3, PO3, PO7, O1, POz, O2, PO8, PO4, P4). Data from the CSP components, single channels or multi-channel clusters were processed with the following procedure. Firstly, for each subject, in each trial pre- and post-stimulation raw data for each channel was bandpass-filtered with a 5-40 Hz 4th order Butterworth zero-phase filter. The data were then detrended (linear trend and mean of data removed) and alpha power was computed. For the POC-region, values were averaged across the nine channels.

### Analysis of Alpha power

We set out to investigate whether tACS applied in the alpha range over visual areas, either in-phase or anti-phase with ongoing visual alpha oscillations, modulates visual alpha oscillations. For this purpose alpha power modulations were analyzed. After the filtering step was performed, we extracted the data to be analyzed according to the chosen fragments (see *Time intervals*) and applied FFT with zero-padding up to 512 samples to obtain smaller frequency bins. Afterwards, we extracted individual alpha power values by averaging power values in the range of IAF – 1 Hz to IAF + 1 Hz (IAF was determined in the pre-experimental resting state EEG measurement, as described above) for pre- and post-stimulation time windows of each trial, and calculated mean values for each stimulation condition. Modulations of alpha power were tested with a repeated measures ANOVA (ANOVA_RM_) comprising the factors TIME (pre- vs post-stimulation), STATE (eyes open vs eyes closed) and STIMULATION (in-phase vs anti-phase) separately for all signal sources: POz data, POC data, CSP(pre) data and CSP(post) data. Results for these four ANOVA_RM_ models were corrected for multiple comparisons by means of Bonferroni correction. In the case of a violation of the homoscedasticity, degrees of freedom were corrected based on the Greenhouse-Geisser correction. Statistical analysis was performed in R (R Core Team [2016]), using the package *afex* (Singmann et al., 2018) running in RStudio (RStudio Team 2016). Generalized Eta Squared (Bakeman 2005) and Cohen’s D (Lakens 2013) served as estimates of effect sizes. Post-hoc contrasts and marginal means (Searle et al., 1980) were calculated via the *emmeans* package.

### Analysis of average power spectra

To investigate modulations of neural oscillations other than in the [IAF-1, IAF+1] frequency band, we computed and analyzed power spectra averaged across all trials for the eyes closed (EC) and eyes opened (EO) conditions for the pre- and post-stimulation data from POz, from CSP and resting state POz data. For the resting state we used the same data as for the estimation of IAF (2 min EC, 2 min EO), but applied a segmentation of 1 s intervals to have the same structure as with the pre- and post-stimulation data and, thereby, obtained two blocks of 50 trials, separately for EO and EC. The spectra were averaged across all segments separately for the EO and EC conditions.

### Analysis of alpha power modulations across the time course of the experiment

As the analysis of alpha power revealed pre- to post-stimulation decreases for the CSP(pre) data (see Results), in an exploratory post-hoc analysis investigated whether alpha power modulations change across the time course of the experiment. For this purpose CSP(pre)-based alpha power values were extracted separately for each block of the experiment, similar to what is described above. For each trial, pre- to post-stimulation power modulations in percent were calculated and averaged separately for each block and experimental condition and statistically tested with an ANOVA_RM_ with the factors BLOCK, STATE and STIMULATION.

### Analysis of alpha power modulations separately for different post-stimulation time windows

In a second post-hoc analysis we wanted to investigate the time scale of post-stimulation alpha power decreases. For this purpose we examined the temporal evolution of the effect by comparing a pre-stimulation time window ([−600 to −100] ms) to four different 500-ms long, overlapping, post-stimulation time windows ([100 to 600] ms, [200 to 700] ms, [300 to 800] ms, and [400 to 900] ms in relation to the onset of stimulation). Pre-to-post modulations of power values for the CSP(pre) data were modeled with an ANOVA_RM_ comprising the factors TIMEBIN, STATE and STIMULATION.

## Results

### No evidence of an alpha power modulation by tACS at single channel level (POz) and in a parieto-occipital cluster (POC)

Mean alpha power for single channel data for exemplary subjects and for the group across the experimental blocks are presented in Figure 2. In general, it shows a diverse pattern: mixed dynamics without a clearly expressed direction of changes for individuals and no significant changes between pre- and post-stimulation on the group level. However, in comparison to average resting state values, there is a slight increase of alpha power for all eyes open blocks and a slight decrease for the first two eyes closed blocks. On the group level alpha power values vary largely, for eyes closed average std = 6.806, for eyes open average std = 3.817. An ANOVA_RM_ revealed alpha amplitude values extracted from POz and a general parieto-occipital cluster to be dependent on the factor STATE (i.e., to be higher when eyes were close as compared to eyes open; *p* < .001), but not to be modulated by tACS or any interaction between the experimental factors (all *p*s > .292) (see Table 1).

**Table 1.** ANOVA_RM_ table representing results of the analysis of FFT-derived power values separately for all signals. Significant effects of are marked by asterisks. ***: *p* < .001, **: *p* < .01. *p*-values are Bonferroni corrected for the four ANOVA_RM_ models.

| signal | factor | df | *F* | *p* | $\eta^2_G$ |
| --- | --- | --- | --- | --- | --- |
| CSP(post) | TIME | (1,19) | 0.524 | 1 | 0 |
|  | **STATE\*\*\*** | **(1,19)** | **39.322** | **< .001** | **0.037** |
|  | STIMULATION | (1,19) | 0.111 | 1 | 0 |
|  | TIME X STATE | (1,19) | 0.647 | 1 | 0 |
|  | TIME X STIMULATION | (1,19) | 0.025 | 1 | 0 |
|  | STATE X STIMULATION | (1,19) | 1.636 | 0.8651 | 0 |
|  | TIME X STATE X STIMULATION | (1,19) | 0.709 | 1 | 0 |
| CSP(pre) | **TIME\*\*** | **(1,19)** | **17.14** | **0.0022** | **0.004** |
|  | **STATE\*\*\*** | **(1,19)** | **26.6** | **< .001** | **0.09** |
|  | STIMULATION | (1,19) | 0 | 1 | 0 |
|  | TIME X STATE | (1,19) | 7.246 | 0.0578 | 0.001 |
|  | TIME X STIMULATION | (1,19) | 0.224 | 1 | 0 |
|  | STATE X STIMULATION | (1,19) | 1.114 | 1 | 0 |
|  | TIME X STATE X STIMULATION | (1,19) | 0.712 | 1 | 0 |
| POC | TIME | (1,19) | 3.684 | 0.797 | 0.002 |
|  | **STATE\*\*\*** | **(1,19)** | **20.38** | **< .001** | **0.12** |
|  | STIMULATION | (1,19) | 1.53 | 0.4588 | 0.001 |
|  | TIME X STATE | (1,19) | 1.725 | 1 | 0 |
|  | TIME X STIMULATION | (1,19) | 1.544 | 0.876 | 0.001 |
|  | STATE X STIMULATION | (1,19) | 0.432 | 1 | 0 |
|  | TIME X STATE X STIMULATION | (1,19) | 0.36 | 1 | 0 |
| POz | TIME | (1,19) | 1.3 | 1 | 0 |
|  | **STATE\*\*\*** | **(1,19)** | **43.192** | **< .001** | **0.253** |
|  | STIMULATION | (1,19) | 3.923 | 0.2492 | 0 |
|  | TIME X STATE | (1,19) | 0.347 | 1 | 0 |
|  | TIME X STIMULATION | (1,19) | 0.015 | 1 | 0 |
|  | STATE X STIMULATION | (1,19) | 0.226 | 1 | 0 |
|  | TIME X STATE X STIMULATION | (1,19) | 0.166 | 1 | 0 |

**Figure 2.**
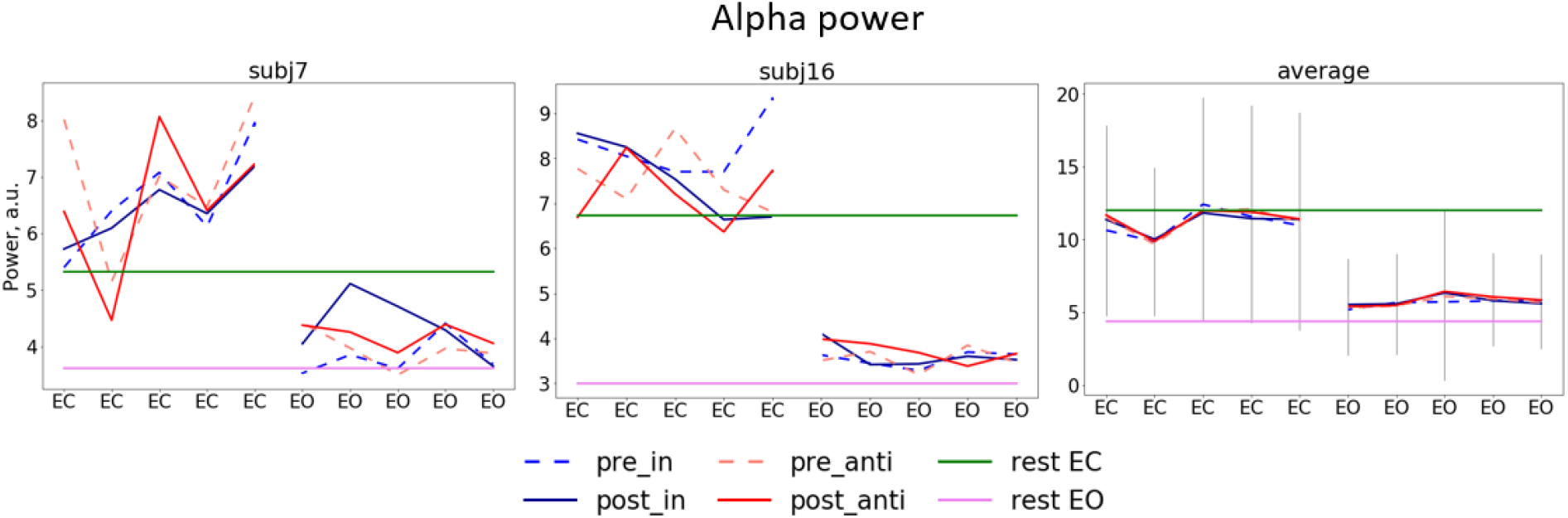
Dynamics of alpha power separate for stimulation blocks. Mean pre/post stimulation values for in-phase and anti-phase relation (pre_in, post_in, pre_anti, post_anti) for exemplary subjects (the left and the center plots) and for the group average (the right plot) 1^st^ 5 blocks (EC) correspond to blocks in which participants had their eyes closed and 2^nd^ 5 blocks (EO) eyes open, green and pink horizontal lines represent alpha power for eyes closed and eyes opened in resting state, gray lines represent average standard deviation.

### Spatially-specific and individualized analysis on the basis of CSP components

Spatial topographies for exemplary subjects and average spatial topographies for CSP(pre) and CSP(post) are shown in Figure 3. Patterns show weights of the inverse matrix of CSP filters. Because the polarity of the distribution is arbitrary, the topographies were normalized by changing the sign to positive in case of negative weights for the patterns at the occipital-parietal area. For the estimation of the average topographies, the patterns of each subject were first normalized by the standard deviation of all channel weights. These normalized patterns could then be averaged. While individual topographies showed some variations in the precise distribution, the average topographies clearly represented the occipito area, which was our targeted region. In addition, components were constructed with narrow band-pass filtering (7–14 Hz), thus, the data obtained with CSP components mainly extracts alpha oscillations.

**Figure 3.**
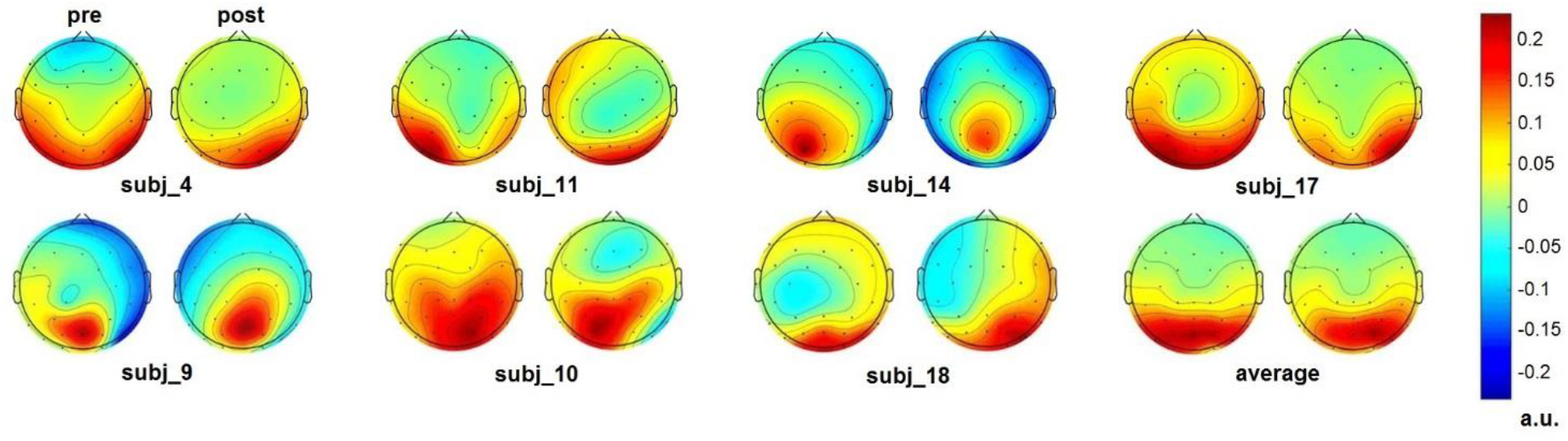
Single subject and average CSP topographies. For each subject as well as the average of all subject topographical distributions of the inverse matrix of the CSP filters are shown for CSP(pre) on the left and CSP(post) on the right.

### Alpha power is modulated by tACS in individual spatial components

Alpha power values can be seen in Figure 4A separately for the experimental conditions, the analyzed electrodes, and CSP-components. As visible in Figure 4A and 4B, pre- and post-stimulation values only differ for power values derived from the CSP(pre) data. This difference is substantiated by the main effect for the factor time (*p* = .002; see Table 1): Across experimental conditions pre-stimulation power values are larger (*M* = 3.23; *CI* = [2.48, 3.67]) than post-stimulation power values (*M* = 3.08; *CI* = [2.64, 3.82]). Additionally, there is a trend for an interaction of factors TIME X STATE (*p* = .058), with larger pre- to post-stimulation decreases when eyes were closed (*M* = −0.23; *SE* = 0.047) as compared to eyes open (*M* = −0.076; *SE* = 0.047). As revealed by the main effects STATE, for all signals (*p*s < .001; see Table 1) alpha power values were always larger when eyes were closed as compared to when they were open.

**Figure 4.**
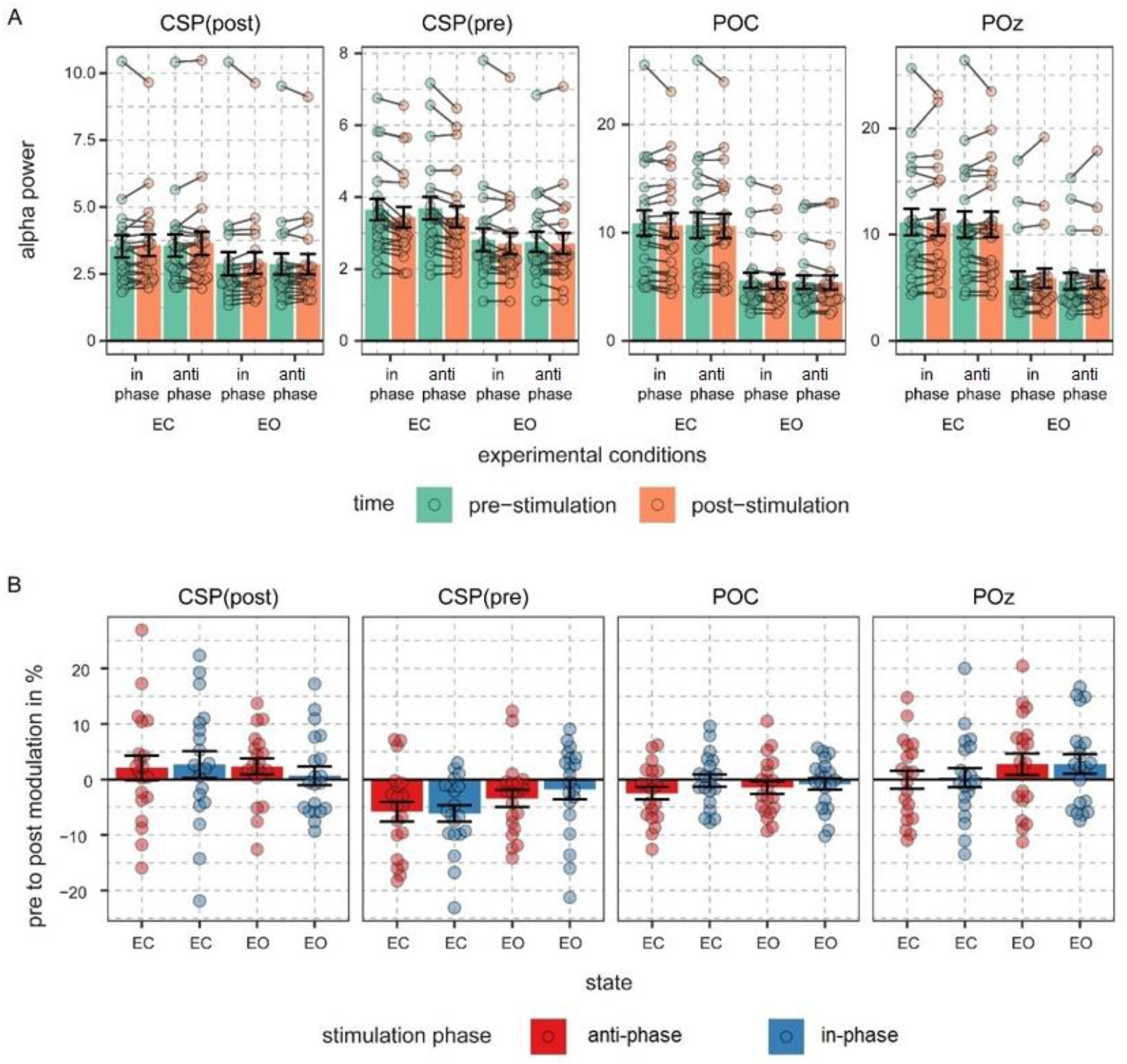
Alpha power values and pre- to post-stimulation modulations of alpha power. A) FFT-derived power values were calculated for a 500-ms long pre- and post-stimulation time window separately for all experimental conditions and signal sources. Note different scales. Connected dots represent single subjects’ pre- to post-alpha power changes. B) Pre- to post-stimulation modulations in % are shown for all signals and conditions. Dots represent single subject data and error bars represent standard error of the mean.

### Modulation of power of neural oscillations is specific to the alpha range

Mean spectra averaged across all trials for different conditions, stimulation relations, with data from POz, both CSP components, parietal-occipital cluster (POC), as well as relations of post- to pre-stimulation are shown in Figure 5. The analysis of the spectra revealed visible differences between pre- and post-stimulation time for signals extracted from both CSP components (Figure 5A) for when participants had their eyes closed as well as open. As visible in Figure 5B, these differences have their maxima in the alpha range without prominent changes to frequencies other than in a range near individual alpha bands.

**Figure 5.**
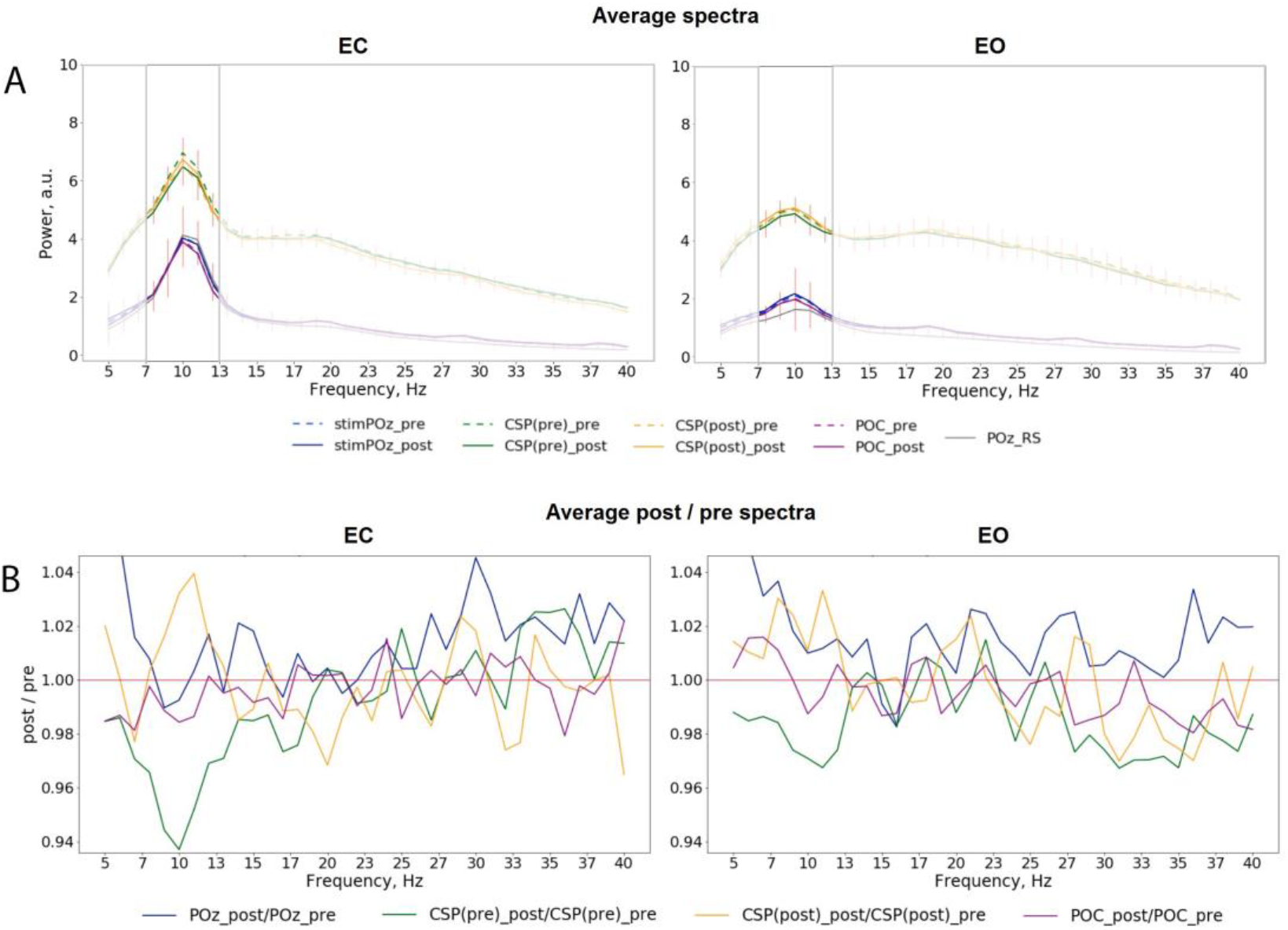
A) Average pre- and post-stimulation spectra for different eye conditions, including both in- and anti-phase trials: a) eyes closed (left), eyes opened (right), red vertical lines represent standard deviation of average difference between pre- and post-stimulation B) Average relation of post-/pre-stimulation spectra

### Modulations of alpha power in a spatial region does not change across the time course of the experiment

In an exploratory post-hoc analysis, we investigated whether the modulation of CSP(pre)-derived alpha power values by tACS changed across the experimental blocks. As visible in Figure 6, there was no systematic change of pre- to post alpha-power modulations across the time course of the experiment. Consequently, modeling CSP(pre)-derived pre- to post power modulations with an ANOVA_RM_ model revealed the factor BLOCK to be insignificant (see Table 2). Therefore, overall pre- to post-stimulation modulations of alpha power seemed to be stable across the experiment. Pre- to post-stimulation decreases were, however, dependent on the state, as revealed by the main effect for the factor STATE (*p* = .031) and was larger when eyes were closed (*M* = −5.786; *CI* = [−8.756, - 2.816]) as compared to eyes open (*M* = --1.829; *CI* = [−4.8, 1.141]).

**Table 2.** ANOVA_RM_ table representing results of the analysis of FFT-derived pre-to post-stimulation power modulation for the CSP(pre) data for time course analysis of the stimulation effects across the experiment. Significant effects of are marked by asterisks. *: *p* < .05.

| factor | df | F | p | $\eta^2$ |
| --- | --- | --- | --- | --- |
| **STATE\*** | **(1,17)** | **5.563** | **0.0306** | **0.022** |
| STIMULATION | (1,17) | 0.684 | 0.4196 | 0.002 |
| BLOCK | (3.13, 53.2) | 2.009 | 0.1213 | 0.021 |
| STATE X STIMULATION | (1,17) | 0.279 | 0.604 | 0.001 |
| STATE X BLOCK | (3.33, 56.6) | 0.289 | 0.8522 | 0.003 |
| STIMULATION X BLOCK | (3.58, 60.9) | 0.43 | 0.766 | 0.004 |
| STATE X STIMULATION X BLOCK | (3.22, 54.7) | 1.571 | 0.2043 | 0.017 |

**Figure 6.**
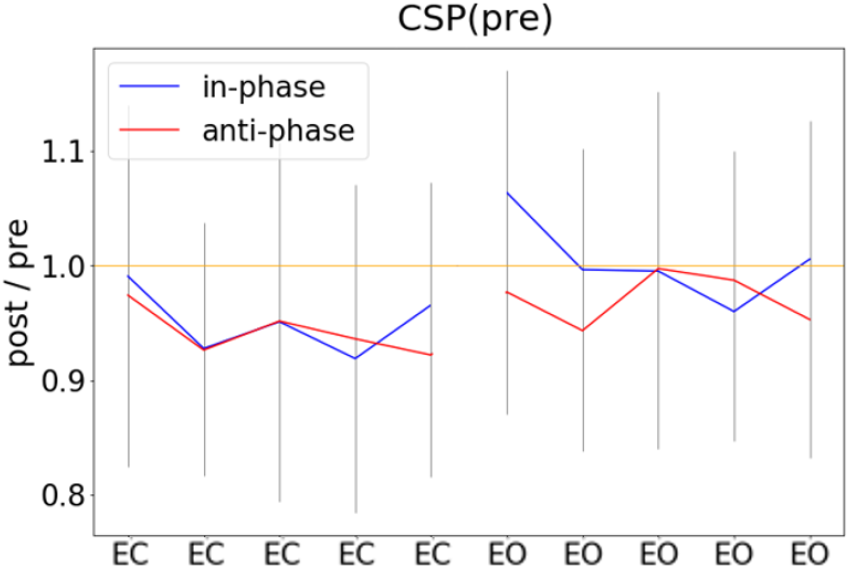
Pre- to post-stimulation power modulations computed separately for experimental blocks. Pre- and post-stimulation ratios of FFT-derived power values were calculated for a 500-ms long pre- and post-stimulation time window separately for all experimental conditions and experimental blocks, gray lines represent standard deviation.

### Modulation of alpha power in a spatial region is transient

We investigated the time scale of post-stimulation alpha power decreases by additionally modeling the effects of pre- to post-stimulation power modulations separately for different overlapping post-cue time windows. The analysis revealed the factor TIMEBIN to be significant (*p* = .018; see Table 3). Post-hoc linear contrasts revealed pre- to post-stimulation decreases to be modeled best by a linear decrease (*t*(57) = 3.497; *p* = .001). Overall, independent of the experimental condition (i.e., eyes open vs. closed; stimulation in- vs. anti-phase) the pre-to post-stimulation decreases are largest right after the stimulation and then decay across subsequent time windows (see Figure 7).

**Table 3.** ANOVA_RM_ table representing results of the analysis of FFT-derived pre-to post-stimulation power modulation for the CSP(pre) data for analyzing the time scale of the stimulation effects. Significant effects of are marked by asterisks. *: *p* < .05.

| factor | df | *F* | *p* | $\eta^2$ |
| --- | --- | --- | --- | --- |
| STATE | (1,19) | 3.201 | 0.0895 | 0.037 |
| STIMULATION | (1,19) | 0.58 | 0.4557 | 0.004 |
| **TIME BIN\*** | **(1.95, 37)** | **4.537** | **0.018** | **0.006** |
| STATE X STIMULATION | (1,19) | 0.76 | 0.3941 | 0.007 |
| STATE X TIME BIN | (1.34, 25.4) | 1.303 | 0.2772 | 0.002 |
| STIMULATION X BLOCK | (1.41, 26.9) | 0.055 | 0.8926 | 0 |
| STATE X STIMULATION X BLOCK | (1.8, 34.2) | 0.141 | 0.8487 | 0 |

**Figure 7.**
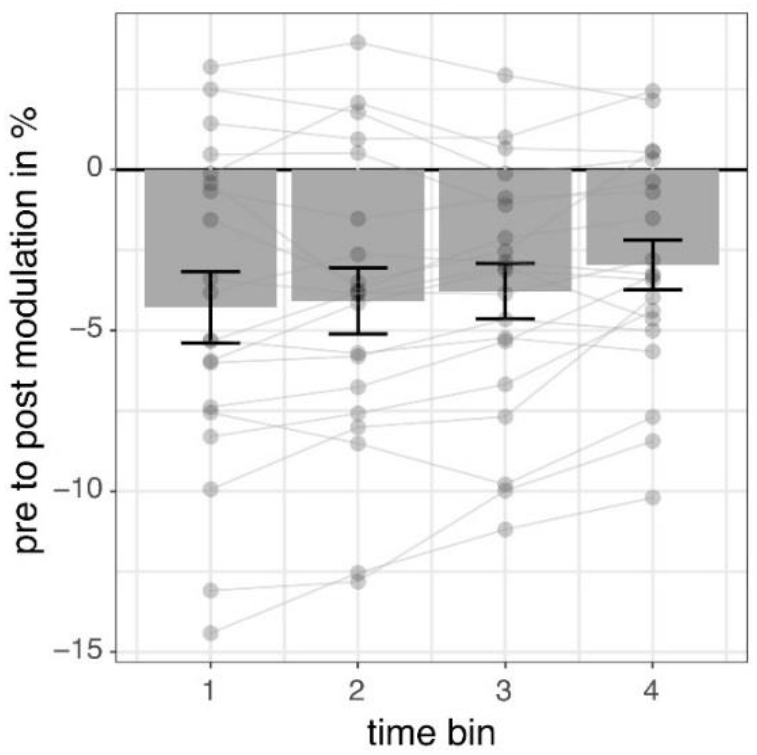
Pre- to post-stimulation power modulations separately for different time windows. The data represents pre- to post-stimulation modulations of alpha power from a 500-ms long pre-stimulation time window to four different 500-ms long overlapping post-stimulation time windows averaged for all experimental conditions in per cent. Post-stimulation time bins are made up by data from the following time windows: [100; 600] ms, [200; 700] ms, [300; 800] ms, [400; 900] ms in relation to onset of the stimulation and always compared to the same pre-stimulation time window [−600; −100] ms. Error bars represent standard error of the mean and dots represent single subjects.

## Discussion

The aim of our study was to investigate effects of tACS applied bilaterally over the visual cortex, tuned to neural alpha oscillations with a closed-loop EEG-tACS setup on visual alpha oscillations. Specifically, we have studied stimulation effects of tACS applied either in-phase or anti-phase with ongoing alpha oscillations during periods of a high-amplitude vs. low-amplitude alpha oscillations on the amplitude of alpha oscillations.

Overall, we found a decrease in alpha amplitude immediately after tACS when accounting for individually spatially specific alpha components with a cross-validation procedure. While these changes had an overall topographical center of gravity in occipital regions, they were individually specific and effects were not observable when data was extracted from a single occipital electrode or a general occipital electrode cluster as in a conventional analysis approach. Although the decreases in amplitude found for a 500-ms long time window seem to be only transient and attenuate across the range of 400 ms, they were constant across the time course of the experiment.

In contrast to previous studies, we found a decrease of alpha amplitude as a response to tACS. Zaehle and colleagues (2010) previously reported an increase of alpha amplitude after 10 min of tACS applied over occipital areas at the individual alpha frequency. Similarly, in various subsequently published studies the application of alpha tACS over visual areas in the range of minutes led to an increase in alpha amplitude (Neuling et al., 2013; Helfrich et al., 2014; Kasten et al., 2016). While these studies differ in their overall stimulation duration, some studies used intermittent short stimulation protocols closer to the design in our study. Strüber et al. (2015) used an experimental protocol similar to ours by intermittently applying 1-s long stimulation trials using conventional tACS. They found no evidence of a modulation of alpha power by tACS. However, they analyzed data only from a single channel (POz) and a longer time interval (1 s). Our data revealed significant stimulation effects to be present only in individual spatial components, but absent at POz and to be transient as they decreased within 400 ms after the end of stimulation. Vossen et al. (2015) applied longer stimulation durations with a different stimulation electrode montage (bilaterally over PO7/PO9 and PO8/PO10) and found that only 8-s intermittent stimulation, but not 3 s, led to pronounced alpha amplitude increases. In another study, Silva et al. (2018) applied intermittent non-adaptive stimulation of a 6-s duration to investigate the influence of tACS on somatosensory perception and found that such stimulation was not sufficient to induce significant causal effects on EEG-measured alpha oscillations.

Because stimulation protocols with longer stimulation durations seem to lead to an increase in alpha amplitude and studies employing trains of stimulation with a duration of 3 s or less either found no evidence for a stimulation effect or a decrease in amplitude, as we did here, it is tempting to speculate about the parameters that shape the stimulation effect. If decreases and increases in amplitude represent two extreme cases, is there a stimulation duration that represents a transition from one to another? What additional factors may contribute to shaping the stimulation effects? Recent studies suggest that the brain state plays a crucial role: When eyes were closed or the room was not illuminated, alpha tACS did not lead to an increase in the amplitude, suggesting that tACS may not modulate the amplitude of oscillations that are already in a high amplitude state (Neuling et al., 2013; Ruhnau et al., 2016). An additional factor may be related to the electrode positioning. When in anti-phase and inter-hemispherically stimulating two coupled mu-alpha generators in the somatosensory system, we previously found a decrease of mu-alpha amplitude after a 5-min tACS application (Gundlach et al., 2017). This decrease was also found for inter-hemispheric tACS targeting theta oscillations (Garside et al., 2015). In a computational study, simulating stimulation after-effects in neural networks with nodes coupled with a time delay in-phasic stimulation led to amplitude increases, while anti-phasic stimulation led to no increases in oscillatory activity (Kutchko & Fröhlich, 2013). Therefore, further studies parametrically manipulating different factors such as duration and electrode position are required to map the effects of tACS more completely.

We found stimulation related decreases in the amplitude to be independent of the phase relationship between ongoing alpha oscillations and the tACS signal. Both in-phase as well as anti-phase stimulation (i.e., stimulation phase being identical to phase of ongoing alpha oscillation measured over POz vs. shifted by 180 degrees and thereby reversed in polarity) disrupted ongoing alpha oscillations. This finding is difficult to reconcile with a stimulation effect mediated by entrainment. While in animal studies online effects were found to be directly related to an entrainment of ongoing neural activity by the applied electric oscillation (Ozen et al., 2010; Reato et al., 2010; Fröhlich & McCormick, 2010), in human studies investigating post-stimulation modulations of oscillations it was proposed that these offline effects may stem from LTP/LTD related effects (Vossen et al., 2015; Zaehle et al., 2010; Vosskuhl et al., 2018). Like others, we previously reported on tACS driven decreases in the amplitude of ongoing oscillations (Garside et al., 2015; Gundlach et al., 2017). While stimulation locations and protocols differed in these studies, the findings common to them and reported here showed, that amplitude decreases even beyond the stimulation period cannot be caused and explained by a mere entrainment of ongoing oscillations by tACS on its own. Thus, it seems to be the case that offline effects are caused by neurophysiological mechanisms different from entrainment. Interestingly, similar to the effects found in the animal studies described above, online effects of tACS measured in humans *during* the stimulation are consistent with an entrainment of neural activity by tACS. For instance, behavioral modulations depend on the stimulation frequency (Joundi et al., 2012; Santarnecchi et al., 2013) and phase of the tACS signal (Neuling et al., 2012; Gundlach et al., 2016). This suggests that tACS effects may be caused by two different and potentially distinct mechanisms (see Heise et al., 2019): tACS may lead to an online entrainment of ongoing oscillations and additional changes in neural plasticity responsible for stimulation outlasting offline effects. While Kar and Krekelberg (2014) found tACS-induced changes in neural adaptation to be potentially closely linked to changes in neural plasticity, the functional underpinnings of such effects as well as the relationship between online entrainment and offline neural plasticity remain unknown.

Interestingly, the stimulation effects in our study seem to be only transient, as they decreased after the end of the stimulation and were not different across the time course of the experiment (i.e., there was no evidence for a cumulative build-up). Effects related to neuroplastic changes seem to depend on the stimulation duration. For instance, neural excitability is longer modulated the longer tDCS was applied (Nitsche et al., 2003), and after-effects of tACS on behavior increased across the time course of the experiment (Heise et al., 2019). A tentative and an alternative interpretation of our findings showing the stimulation effect to be independent of the time in the experiment, could be that the application of short stimulation periods in our experiment did not lead to plastic modulation of alpha generators, but instead briefly disturbed ongoing oscillations. Because alpha rhythm seems to fluctuate between different states of activity level (Freyer et al., 2009; 2011), the transient decrease in amplitude after the application of tACS may index a brief shift of alpha activity towards a lower activity level by tACS. However, when assuming an online entrainment of alpha activity by tACS, it is puzzling that both in- and anti-phase stimulation lead to a similar effect of amplitude attenuation. Specifically, one would hypothesize a synchronous application (no phase difference and the same polarity) not to disrupt the specific oscillation, while an asynchronous application would be more likely to be disruptive. We extensively tested our setup and phase extraction as well as forecast algorithms to ensure that the relation of the stimulation phase and phase of ongoing alpha oscillations was captured correctly (see Zarubin et al., 2018). While small deviations in the phase relation may arise from the phase estimation, forecast process, and underlying assumptions of stationarity, the overall phase relation is thus estimated accurately. If in- and anti-phase stimulation are indeed different in their phase relationship, why would they lead to a similar effect, namely the decrease in alpha amplitude? Several recent studies have suggested that alpha oscillations measured on a macroscale are the product of different alpha generators with different spatial, laminar, or functional profiles rather than being produced by a single generator (Barzegaran et al., 2018; Benwell et al., 2019; Bollimunta et al., 2008; Haegens et al., 2015; Keitel & Gross, 2016; Scheeringa et al., 2016; Schaworonkow and Nikulin, 2019). The potential interaction, coupling, and interdependence of different alpha generators is, however, vastly unknown. There is some evidence that different alpha generators may be antagonistically coupled, as for instance seen in the relationship of visual alpha and sensorimotor mu-alpha (Neuper & Pfurtscheller, 2001; Gerloff et al., 1998) or the focal down- and surround up-regulation of alpha generators in the somatomotor system (Suffczynski, 1999) or the different alpha profiles in different layers of intracortical measurements (Bollimunta et al., 2008, 2011). If different alpha generators were indeed negatively coupled, the up-modulation in one (i.e., by NIBS) could lead to a down-modulation of the other. Because the single trial, cross-validated CSP filtering procedure for each trial and subject extracts components, which either maximize the post-over the pre-stimulation alpha power (or vice versa), this procedure may extract the individual spatially distinct alpha component that shows a decrease in power. This worked better for extracting components that showed a decrease after stimulation, but not an increase after stimulation, suggesting that the disturbing effect for occipital alpha generators may be much more robust.

To our knowledge, we are the first to utilize CSP filtering for the analysis of tACS effects and thus to obtain neural activity selectively tuned to show a decrease or an increase of alpha oscillations following tACS. Although CSP is mainly applied in BCI paradigms to differentiate particular activation patterns, which usually correspond to different anatomical regions (e.g., left or right motor imaginary corresponding to the activation of the right or left motor cortex), this method also can be useful for discriminating between activity in the same spatial region, which was previously shown for alpha oscillations with standard and deviant visual stimuli (Tugin et al., 2016). In our study CSP was used to provide spatial filters maximally discriminating activity between the periods with and without stimulation. This in turn allowed the contribution of stimulation-related neural changes to be maximized, while attenuating irrelevant neural activity typically masking effects of stimulation in the sensor space. As previously mentioned, tACS always affects a broad region of neural populations, and thus studying its influence based only on data from a single or few channels is only an approximate simplification. Such approaches lead to a significant reduction of the observation space, the omission of region-specific dynamics, and raises the impact of noise and volume conduction in the data. Therefore, to perform a deeper, more comprehensive and more extensive investigation of the respective research question, CSP and other spatial filtering methods should be considered for the analysis of tACS effects and brain stimulation effects in general. Importantly, the modulatory effects of tACS were primarily limited to the occipito-parietal regions – areas targeted with tACS in our study. Such spatial distribution of the observed effects challenges the possibility that the effects of tACS might have been due to the stimulation of the scalp (Voroslakos et al., 2018; Asamoah et al., 2019). In this case, we would expect attenuation of alpha/mu oscillations over the sensorimotor areas, which was not the case here.

One limitation of our study is that our design may be suboptimal in promoting plastic changes of neural activity developing during the time of the stimulation. Using a purely event-related design, the stimulation phase varied randomly between in-phase and anti-phase with ongoing alpha oscillations. If tACS were able to lead to an online entrainment of ongoing neural oscillations as a potential prerequisite for offline effects, one could hypothesize that this effect would be more pronounced when the same phase-orientation (“in” or “anti”) were utilized over the whole duration of the stimulation block or experiment (Deans et al., 2007; Fröhlich & McCormick, 2010; Ozen et al., 2010; Helfrich et al., 2014). Varying the phase relationship across trials might have interfered with an effect that relies on accumulating over time and may have masked the potential effects of tACS.

Closed-loop tACS and, in general, adaptive NIBS in comparison to conventional stimulation protocols have several advantages, which could be potentially beneficial for the whole brain stimulation research field and transform it to a more reliable and clinically applicable approach. As mentioned by Zrenner et al. (2016), these advantages include: personalized neuromodulation to decrease inter-individual variability of effects, analysis of network reorganization dynamics, such as during stroke, for instance, to aid rehabilitation and target as well as specifically modify potentially different plasticity patterns. One of the main challenges impeding the development of closed-loop tACS, however, is the fact that the analysis of online effects during stimulation is compromised by massive stimulation-induced artifacts. In principle, the signal to be analyzed and modulated (e.g., a signature of alpha oscillations measured in the EEG) is overwritten by the tACS signal, which is several magnitudes larger but covers the same spatial and temporal space. Thus, substantial efforts in recent studies have been directed to the development of artifact elimination methods using different techniques and experimental protocols (Witkowski et al., 2015; Kohli & Casson, 2019; Kasten et al., 2018; Noury & Siegel, 2017). However, intermittent stimulation allows one to follow another approach while still based on adaptive principles. By studying the immediate after-effects in intervals between periods of stimulation without artifacts, such studies may contribute towards exploring related online effects and enhance the understanding of tACS effects and mechanisms in general. One prominent recent study with phased-locked closed-loop stimulation was presented by Mansouri et al. (2018). By using intermittent stimulation with very short (5 ms) square-wave pulses and an artifact removal procedure using a spline interpolation (Waddell et al., 2009), they were able to extract the artifact-free EEG signal and thus control for the actual phase locking of delivered stimulation and ongoing oscillation in alpha and theta bands. Such approaches may help to elucidate the role of adaptive NIBS on brain activity and ultimately the role of brain activity on cognition, perception, and behavior in general.

In summary (see Figure 8), we found that short-time intermittent tACS applied over occipital regions (Cz and Oz), as used in previous studies, induces a transient suppression of occipital alpha generators, leading to a decrease in alpha power in spatially specific components centered over a parietal-occipital region. This effect was independent of the phase relationship between the tACS signal and alpha oscillations. This suggests to us that these offline effects of short-timed intermittent tACS are not explainable by entrainment alone but rather require neuroplastic changes or a transient disruption of neural oscillations. These effects were only visible in individual spatial alpha components, but not in a broad occipital cluster or pre-selected electrode. Our study thus supports the notion that the response to tACS differs inter-individually and that even intra-individual effects are shaped by the interplay between different alpha generators. This favors stimulation protocols as well as analysis regimes exploiting inter-individual differences to more efficiently induce as well as more reliably reveal otherwise hidden stimulation effects and thereby comprehensively study the effects and the underlying mechanisms of tACS.

**Figure 8.**
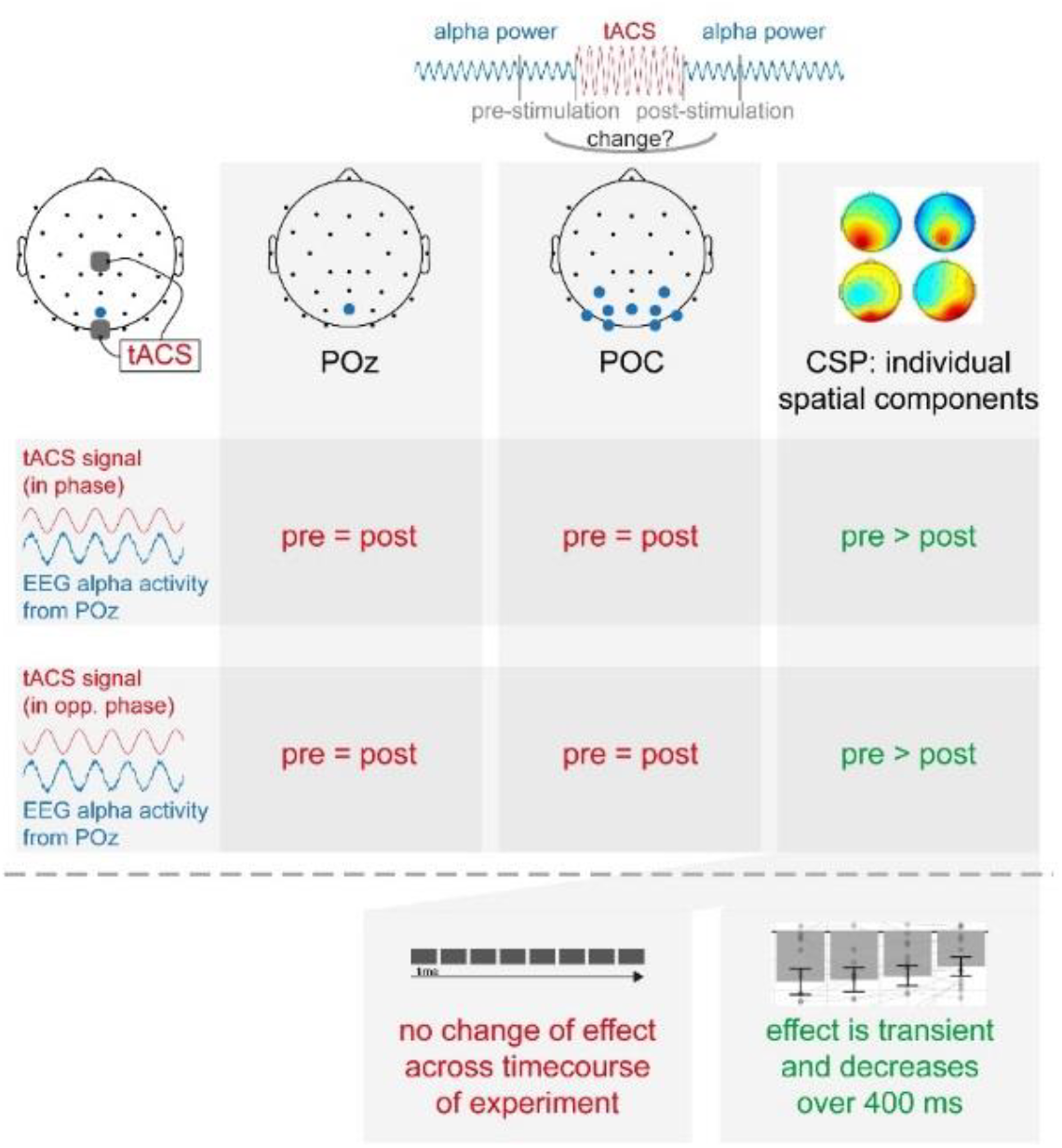
Summary of main experimental findings

